# Combining whole genome shotgun sequencing and rDNA amplicon analyses to improve detection of microbe-microbe interaction networks in plant leaves

**DOI:** 10.1101/823492

**Authors:** Julian Regalado, Derek S. Lundberg, Oliver Deusch, Sonja Kersten, Talia Karasov, Karin Poersch, Gautam Shirsekar, Detlef Weigel

## Abstract

Microorganisms from all domains of life establish associations with plants. Although some harm the plant, others antagonize pathogens or prime the plant immune system, acquire nutrients, tune plant hormone levels, or perform additional services. Most culture-independent plant microbiome research has focused on amplicon sequencing of 16S rDNA and/or the internal transcribed spacer (ITS) of rDNA loci, but the decreasing cost of high-throughput sequencing has made shotgun metagenome sequencing increasingly accessible. Here, we describe shotgun sequencing of 275 wild *Arabidopsis thaliana* leaf microbiomes from southwest Germany, with additional bacterial 16S rDNA and eukaryotic ITS1 amplicon data from 176 of these samples. The shotgun data were dominated by bacterial sequences, with eukaryotes contributing only a minority of reads. For shotgun and amplicon data, microbial membership showed weak associations with both site of origin and plant genotype, both of which were highly confounded in this dataset. There was large variation among microbiomes, with one extreme comprising samples of low complexity and a high load of microorganisms typical of infected plants, and the other extreme being samples of high complexity and a low microbial load. We use the metagenome data, which captures the ratio of bacterial to plant DNA in leaves of wild plants, to scale the 16S rDNA amplicon data such that they reflect absolute bacterial abundance. We show that this cost-effective hybrid strategy overcomes compositionality problems in amplicon data and leads to fundamentally different conclusions about microbiome community assembly.

## Introduction

Microorganisms affect many important plant traits. Opportunistic microbes can slow down growth or kill a plant, while beneficial ones can prime the plant immune system [1], directly antagonize pathogens [2], or indirectly inhibit pathogens by contributing to a suppressive environment [3]. Microbes may adjust plant hormone levels [4] and participate in nutrient acquisition [5,6], among other processes [7]. Research in this area has revealed that most of the organisms present on and in healthy plant leaves are typically bacteria [8]. With the exception of pathogenic strains, fungi, archaea, and protists such as oomycetes generally have lower abundance on leaves than bacteria, and have also received less attention because they are less easily cultured and are genetically more complex. Due to the difficulty of adequately capturing microbial complexity and diversity, we still lack a good understanding of the composition and dynamics of leaf microbial communities, their abundances on the plant, and how they relate to other aspects of the host biology such as genotype, location, or environmental conditions.

Amplicon sequencing, in which a specific locus common to a target group of organisms (usually 16S rDNA for bacteria and ITS for fungi) is amplified and sequenced, has been the tool of choice for revealing the taxonomic composition of a microbiome. Albeit usually very informative, amplicon sequencing relies on oligonucleotide primers that disfavor or exclude some organisms, with the consequence that some taxa are systematically ignored. Not only are there then no quantitative estimates of the excluded taxa, but there is also no information on the absolute abundances of included taxa. Furthermore, one gene cannot reliably predict the other genes and genetic and metabolic functions in a microbe beyond some conserved features [9,10]. Whole metagenome shotgun sequencing of DNA extracts has become an attractive tool for dissecting complex microbial communities, and databases and algorithms that support such efforts are being developed [11,12]. Shotgun sequencing supplies information on the total DNA content of microorganisms as opposed to a specific locus, which in principle enables us to ask many new types of questions [8,13,14].

We used metagenome sequencing to characterize the leaf-associated (phyllosphere) microbiome of 275 wild *Arabidopsis thaliana* individuals from around Tübingen in southwest Germany, at four different timepoints between 2014 and 2016. Of these, we subjected 176 to 16S rDNA and ITS1 sequencing. We achieved low (<100 Mb) to high (>1 Gb) depths of microbe-associated metagenomic sequences per sample, which we mapped to reference databases. Unsurprisingly, the wild *A. thaliana* microbiota was highly variable between individuals, but there were some clear patterns in the dominant microbes; in particular *Pseudomonas* dominated in some sites and *Sphingomonas* in others. We estimated absolute microbial load as the ratio of microbial reads to plant chromosomal reads in each sample, and observed that the microbial load varied across samples from less than 1% to up to 77% of plant reads. After producing a load-corrected table, we often observed that intertaxa abundance correlations changed in sign compared to compositional amplicon data and metagenome data, in which microbial reads had been normalized by total sum scaling. Finally, we document the relative abundance of eukaryotic and archaeal microbes, revealing a small but noteworthy presence of fungi and oomycetes.

## Results

### Sequencing and analysis approach

To obtain an unbiased picture of microbial diversity and microbial load in wild *A. thaliana* plants, we took a metagenomic shotgun sequencing approach to analyze entire wild phyllospheres. Because plant genomic DNA was expected to dominate such samples, we initially attempted to develop a protocol to enrich the microbial component. To this end, we used genomic DNA from sterilely-grown *A. thaliana* plants in excess as bait to remove plant DNA from prepared shotgun libraries. We were unable to find conditions that allowed for consistent, substantial enrichment of microbial sequences (see Supplementary Information for section on Subtractive Hybridization).

We therefore proceeded to shotgun sequence the aerial portions of 275 non-flowering *A. thaliana* individuals from well-characterized locations in southwest Germany [15]. This compilation spanned two different growing seasons (2014/2015 and 2015/2016) with samplings in winter and early spring. Our strategy captured distinct ecological conditions as well as non-overlapping populations across time due to the winter-annual lifestyle of *A. thaliana* in the region (see Methods). Samples were washed three times vigorously with sterile water to remove loosely-adhering dust and dirt particles prior to freezing samples on the same day as the harvest at −80°C. At a later date, total genomic DNA was extracted and converted into barcoded Illumina short-read sequencing libraries. Plants from the five collections were processed in three batches.

To determine host DNA content, quality-filtered sequencing reads were mapped to the *A. thaliana* Col-0 TAIR10 reference genome [16] with *bwa-mem* using standard parameters [17]. Sequences that did not map to the reference genome were then translated in silico in all six reading frames and aligned against NCBI’s non redundant protein database using DIAMOND [18] with standard parameters, and alignments were processed with MEGAN [19,20].

We reasoned that the number of reads from the plant nuclear genome was highly correlated with diploid cell equivalents and thus fresh weight [21,22]. We therefore scaled the non-plant read counts to the number of reads that could be mapped to any of the five *A. thaliana* chromosomes. This use of plant chromosomal DNA is analogous to studies that use internal ‘spike-in’ controls calibrated to sample weight or volume [23–25]; our ‘spike-in’ is inherent to our samples (Supplementary Figure 1).

About half of non-plant reads in each sample could be assigned to microbial taxa. That the number of non-classifiable reads was positively correlated with the number of microbial reads suggested that these unclassified reads were mostly from portions of microbial genomes not present in the NCBI nr database, rather than being plant sequences not found in the *A. thaliana* reference genome (Supplementary Figure 2). To gauge how likely the unclassified reads were to contain sequences missing from the *A. thaliana* reference genome, we mapped these to additional *A. thaliana* genomes assembled from long-read data [26] including five genomes available in house. The number of unclassified reads that mapped to additional plant genomes was unrelated to the quantity of unclassified reads in the sample; even in samples with up to 21% unclassified reads, the fraction of reads that mapped to the additional reference genomes was less than 1% of the total classifiable plant reads (Supplementary Figure 2). In other words, across all samples, only a small but consistent percentage of unclassified reads was likely to come from the plant. The rest most likely reflects additional microbial sequences. These sequences may belong to noncoding regions or genes from known taxa that have not been assembled and incorporated into the database. Currently, we cannot easily know how many of the nonclassified, but putative microbial reads reflect highly variable sequences of accessory genomes from known taxa, nor how many reflect the presence of microbial taxa that have not yet had their genomes sequenced. Overall, our results were reminiscent of efforts to classify metagenomic reads from soil and human gut, where more than 50% of reads could not be annotated against known databases [27–29].

To further investigate species-level identification of microorganisms and search for microbial functions, we attempted metagenome assembly of all samples. We assembled short reads with MEGAHIT [30] (meta-sensitive preset parameter), filtered out contigs shorter than 200 bp, and assessed standard assembly metrics such as N50, N90, mean contig length, and total assembly size (Supplementary Figure 3). Additionally, we mapped reads back to their corresponding assemblies to determine what fraction of each library was effectively being incorporated into contigs. The N50 of contigs that were at least 200 bp long ranged from 500 to 800 bp, with the maximum length of individual contigs in the different assemblies ranging from 6 to 12 kb and the sum of their lengths ranging from 4 to 160 Mb. The mapping rate of short reads to their respective assemblies was only 25%. We made a further attempt at assembling reads into contigs using metaSPAdes [31]. We tried assembling individual samples separately as well as pooling them by sampling location to increase coverage. This yielded only modest improvements (Supplementary Figure 4). This difficulty to assemble long contigs was an apparent consequence of high sample diversity (Supplementary Figure 5). It paralleled the limited success in assembling metagenomes of deeply sequenced soil, where despite of having over 300 Gb of data, 80% of sequences could not be assembled because of low coverage of individual taxa [27].

To evaluate the reproducibility of our approach and to infer potential biases, we prepared independent libraries for 42 samples. We used the same DNA input for 18 samples, but we also split the plant material after it had been ground in liquid nitrogen and performed independent DNA extractions for 24 samples (Supplementary Figure 6). The comparison of the microbial components of the resulting sequencing libraries revealed that samples from the same plant grouped together in hierarchical clustering and ordination analyses, both for libraries prepared from the same DNA and libraries prepared from different DNA extractions. The inter-sample distances were similar for both types of replicates.

### Overview of microbial taxa in the metagenomes

Overall, we found large variability in the fraction of assignable microbial reads, ranging from 3% to 45% of total read counts in each sample (Supplementary Figure 7). The vast majority of microbial sequences was identified as bacterial (Supplementary Figure 8, Supplementary Figure 2), representing on average 47 families and an average Shannon Diversity of 25 families (Supplementary Figure 9). Taxonomic composition varied across plants (Figure 1), seasons, and locations (Supplementary Figure 10). Sphingomonadaceae and Pseudomonadaceae consistently ranked as the most abundant bacterial families (Fig 1 b). Despite both being very common, they behaved very differently across samples: Pseudomonadaceae varied greatly in their relative abundance, with a few samples having substantially higher counts relative to the rest, while the fraction of Sphingomonadaceae reads was more even across all samples (Fig. 1b, Fig. 3e, Supplementary Figure 11).

**Figure 1.**
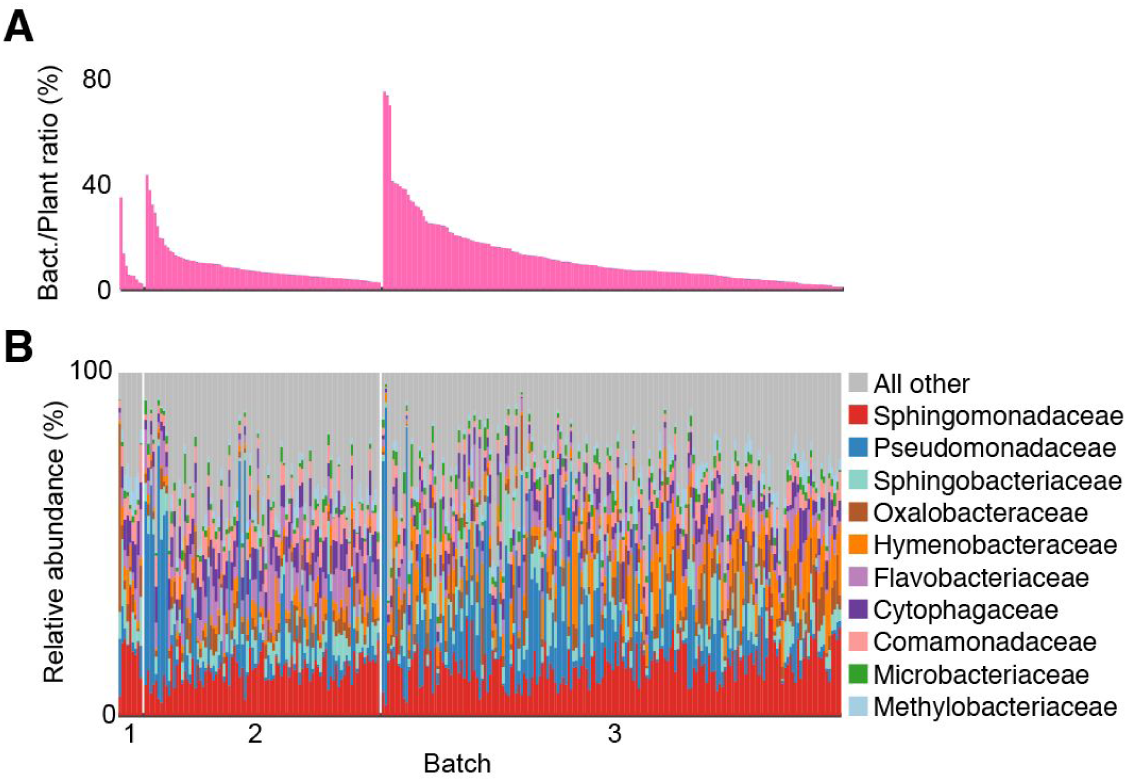
Wild *A. thaliana* leaf microbiomes from shotgun sequencing. **(A)** Relative abundance of bacterial reads as a fraction of total reads in each sample. **(B)** Relative abundance of the 10 most prevalent bacterial families, computed with plant chromosome-scaled read counts. Samples are grouped by processing batch in the same order as in panel (A).

We had chosen an arbitrary depth of sequencing for our effort, and we therefore wanted to learn how much information would be lost by reducing the number of sequencing reads per sample. We made use of the replicated individuals to this end. We downsampled reads from replicated libraries and compared their taxonomic profiles. We estimated that approximately 300,000 non-plant reads constitute a lower bound for robust description of taxonomic profiles (Supplementary Figure 18). This agrees well with similar estimates for human gut microbiome samples [11], and translates into 7.5 million total reads, or just under 1.12 Gb total sequencing reads per sample for 90% of the dataset.

As a counterpoint to downsampling reads, we were curious how much could be gained by having much deeper sequence coverage from a single plant. Therefore, we processed a single plant that was visibly infected with white rust (*Albugo* spp.) and downy mildew (*Hyaloperonospora arabidopsidis*), and that we in addition left unwashed to further potentially increase the fraction of microbial reads. We sequenced this plant to high depth (~20 Gb), which was 5 to 20 fold more coverage than the other samples. Fewer than 40% of reads from this sample mapped to the *A. thaliana* reference genome. Similar to our other samples, about half of the remaining reads could be assigned to microbial taxa, with over 90% coming from bacteria (Supplementary Figure 12). In addition to many *Albugo* spp. and *H. arabidopsidis* reads, we found many of the bacterial taxa already detected in the other samples, and in similar proportions.

### Influence of site, season and host genetics

A common way to compare composition of microbiomes is based on the Bray-Curtis dissimilarity measure. However, a true distance metric is better suited than a dissimilarity measure for other downstream analyses such as principal component analysis [32]. For distance/dissimilarity measures weighted by taxa abundance, highly abundant taxa can strongly skew results while low abundance taxa contribute relatively little information.

To evaluate how the microbiomes of our samples relate to each other, we used, instead of Bray-Curtis dissimilarity, pairwise Euclidean distances and PCA. We first transformed the data by taking the fourth root of the family-level abundance table, including all bacterial, fungal, and oomycete taxa. This transformation corrects for positive skewness in count distribution common in ecological datasets [33], and also decreases the influence of high abundance microbes. No single metadata variable could clearly explain the distributions along the main axes in PCA, although collection site seemed to do best (Figure 2a, 2c, Supplementary Figure 13). Clustering of samples was most clearly driven by the most abundant taxa in each sample, a feature that correlated with collection site. Separation by Pseudomonadaceae or Sphingomonadaceae was apparent when comparing PC1 to PC2, whereas separation by less abundant taxa could be seen when comparing PC2 to PC3 (Supplementary Figure 14).

**Figure 2.**
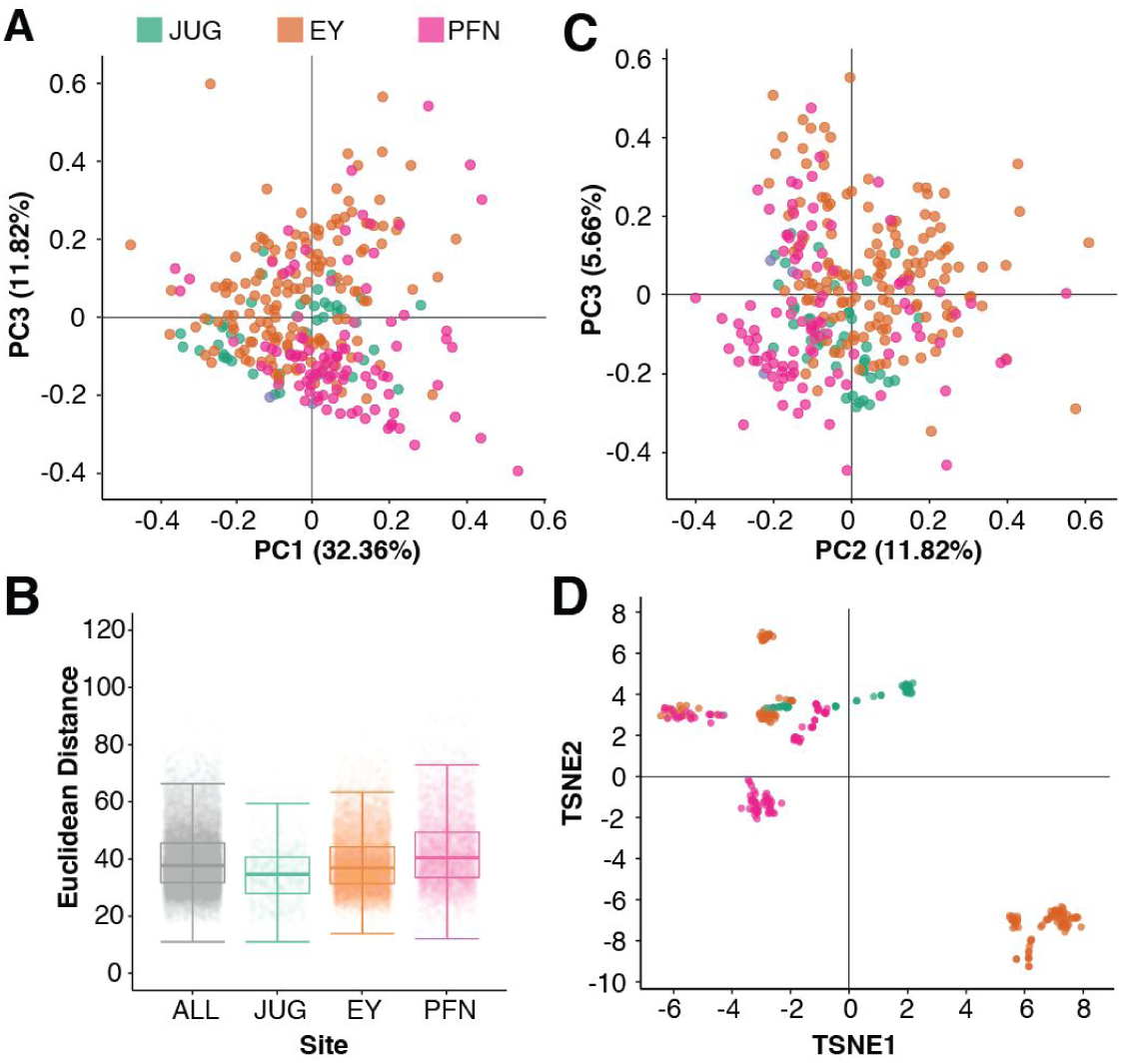
Impact of dominant taxa on microbial community structure. **(A)** Principal component analysis of the fourth root transformed count matrix colored by sampling location. **(B)** Bacterial Euclidean distance distribution across all samples in the dataset (ALL), and from each sampling location (JUG - Kirchentellinsfurt, EY - Eyach, PFN - Pfrondorf). **(C)** t-SNE map of genetic distances (see Methods) with samples colored by location. Distinct genotypes can be identified as clusters of samples; note clear correlation between genetic similarity and sampling site.

Finally, we used the abundant plant reads for host genotyping, using FreeBayes [34] to call close to 1 million SNPs from reads with high quality mapping to the TAIR10 reference genome. There were several clear clusters of plant genotypes (Fig. 2c) correlated with sampling site, in agreement with stands of *A. thaliana* in southwest Germany normally hosting only a limited number of genotypes [15]. It is well known that host genotype can influence the composition of the leaf microbiome [35,36], but as genotype is strongly linked to site in wild populations, both variables are confounded and would require additional data in order to separate the effects of each.

In an orthogonal analysis, we first classified reads broadly as bacteria, fungi, plants, or unclassified, and compared overall sequence similarity between samples in each class using MASH [37], which measures similarities in k-mer abundance. MASH does not consider the taxonomic classification of sequences, but because samples containing different taxa include different sequences and hence different k-mers, this classification-independent analysis captured many of the same patterns in the data. PCoA on MASH distances between bacterial, fungal, plant, or unclassified reads also led to some degree of clustering of samples by collection site (Supplementary Figure 15).

### Intermicrobial correlation networks

Shotgun sequencing provides a minimally-biased estimation of the true abundance of microbes in a microbiome sample. We examined microbial abundances across all samples and under different data transformations to understand colonization patterns and potential intermicrobial relationships. We first made a map of pairwise linear correlations between all bacterial families that passed filtering thresholds (1,000 assigned reads per family in at least 10 samples) using plant-scaled data (equal plant chromosomal reads), relative abundance data, and fourth root of plant-scaled data. Only cooccurrences with a Pearson correlation coefficient greater than | ±0.2 | and with p-value lower than 0.05 after Student’s t-test were used (Supplementary Figure 16). On average, any taxon was positively correlated with 13 other taxa, but this was heavily skewed toward microbes present in many samples, as correlations between taxa only seen in a handful of plants were usually not significant. In addition, bacterial load varied widely across samples and it was positively associated with the abundance of sequences associated with each taxon (Supplementary Figure 17). This resulted in only positive correlations between taxa (figure 3a). That is, a high bacterial load meant a higher abundance of nearly all taxa, although some taxa such as *Pseudomonas* contributed more to microbial load than others. When applying fourth root transformation to the plant-scaled microbial counts, the same trend as in untransformed data was observed for all taxa, the only difference being increased correlation values and an increased number of correlated taxa pairs.

**Figure 3.**
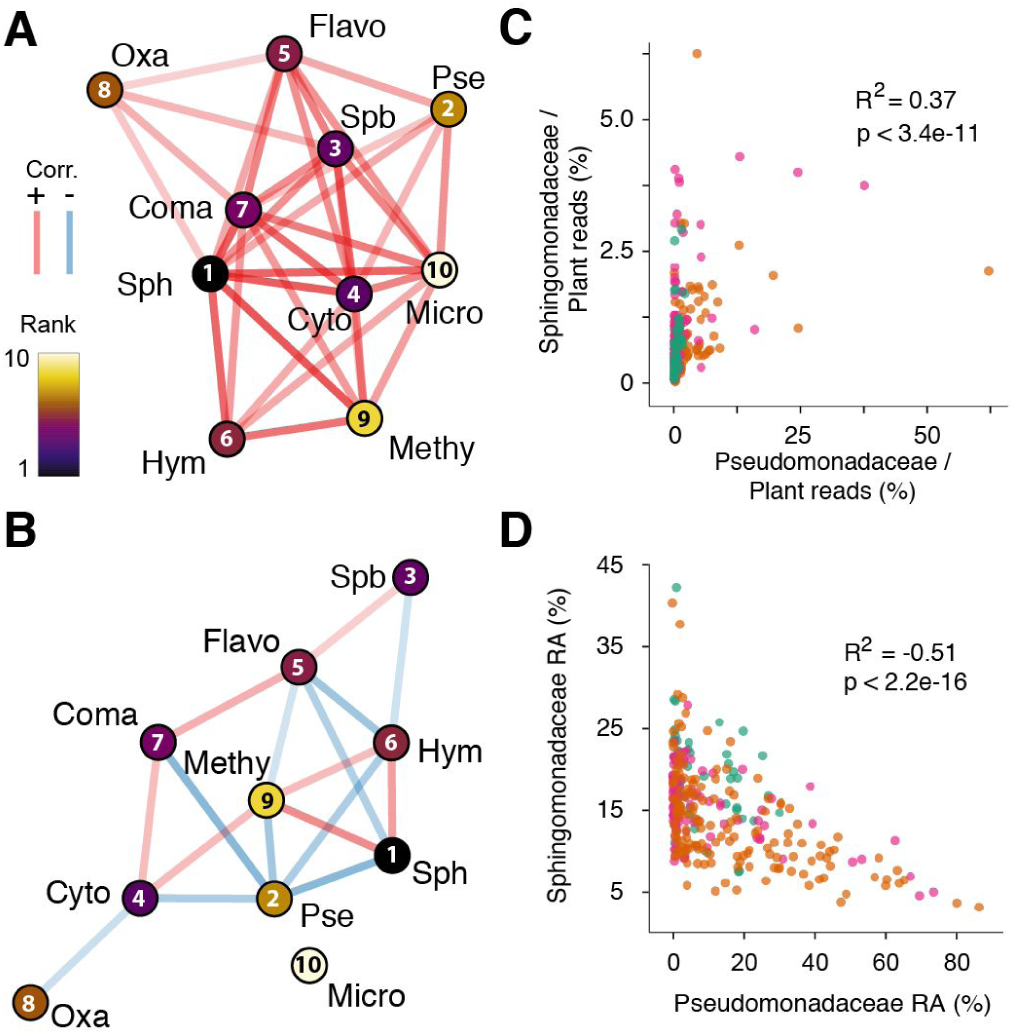
Opposite intertaxa correlations inferred from absolute and relative abundance data. **(A, B)** Correlation networks of 20 most abundant taxa at family level. Nodes represent individual taxa and edges correspond to statistically significant (p < 0.05, R^2^ > 0.2) Pearson correlation between taxa across all samples. Colors indicate direction of correlation (red - positive, blue - negative), transparency reflects correlation strength. Taxa are colored relative to mean rank in the dataset (scale on top). Labels on top of nodes indicate bacterial families as shown in Fig. 2. Pse - Pseudomonadaceae, Sph - Sphingomonadaceae, Flavo - Flavobacteriaceae, Hymn - Hymenobactereaceae, Methy - Methlobacteriaceae, Coma - Comamonadaceae, Cyto- Cytophagaceae, Oxa - Oxalobacteraceae, Micro - Micrococcaceae, Spb - Sphingobacteriaceae. **(A)** Network based on scaled load data. **(B)** Network based on relative abundance data. **(C)** Correlation between plant scaled Sphingomonadaceae and Pseudomonadaceae bacterial load. **(D)** Correlation between relative Sphingomonadaceae and Pseudomonadaceae relative abundance.

If bacterial data in all samples are transformed to relative abundance prior to analysis - a necessity for data without internal standards such as most amplicon data - taxa abundance estimations become constrained because the sum of all taxa is constant, greatly confounding the directionality of their correlation. When such a transformation was applied to our dataset, many of the positive correlations between taxa either disappeared or became negative, including the one between Pseudomonadaceae and Sphingomonadaceae (Figure 3). If we did not have data on bacterial load, it would be tempting to jump on this as evidence for widespread antagonism between these families in the wild. Indeed, antagonism between *Pseudomonas* and *Sphingomonas* is known to occur in laboratory conditions [38]. However, if widespread antagonism exists between members of these families, our data did not reveal it. Spurious conclusions due to compositional data are a common and well-documented problem that several analysis methods, such as the centered-log-ratio transformation, attempt to overcome [39–43].

### Concordance between metagenome and amplicon data

In order to contrast information from metagenome and amplicon sequencing, we focused on the largest batch, batch 3, with 176 samples. We PCR amplified and sequenced the V4 region of bacterial 16S rDNA and the fungal ITS1 region for these samples. Because the V4 16S rDNA sequence of *A. thaliana*-associated cyanobacteria is indistinguishable from that of chloroplasts, reads with cyanobacteria assignments were ignored and cyanobacteria reads were also removed from the metagenome dataset for a fairer comparison. The agreement between assignment of bacterial families based on 16S rDNA amplicons and metagenomes was very high (Fig. 4A), with an overall Pearson coefficient of correlation R^2^ of 0.78 on fourth root transformed data (Figure 4B). Among the top taxa, compared to metagenomics estimates, 16S rDNA estimates were slightly lower for Pseudomonadaceae, and slightly higher for Sphingomonadaceae, Sphingobacteriaceae, and Oxalobacteraceae (Fig. 4A,B). In a complementary comparison, we extracted only the 16S rDNA sequences from the metagenome reads and classified them using *phyloFlash* [44]. When plotted against 16S rDNA amplicons that had been subsampled to match the metagenome 16S rDNA read counts, the overall correlation and overestimation/underestimation trends were the same as for the comparison of amplicons with all metagenome reads (Fig. 4E, compare to Fig. 4B).

**Figure 4.**
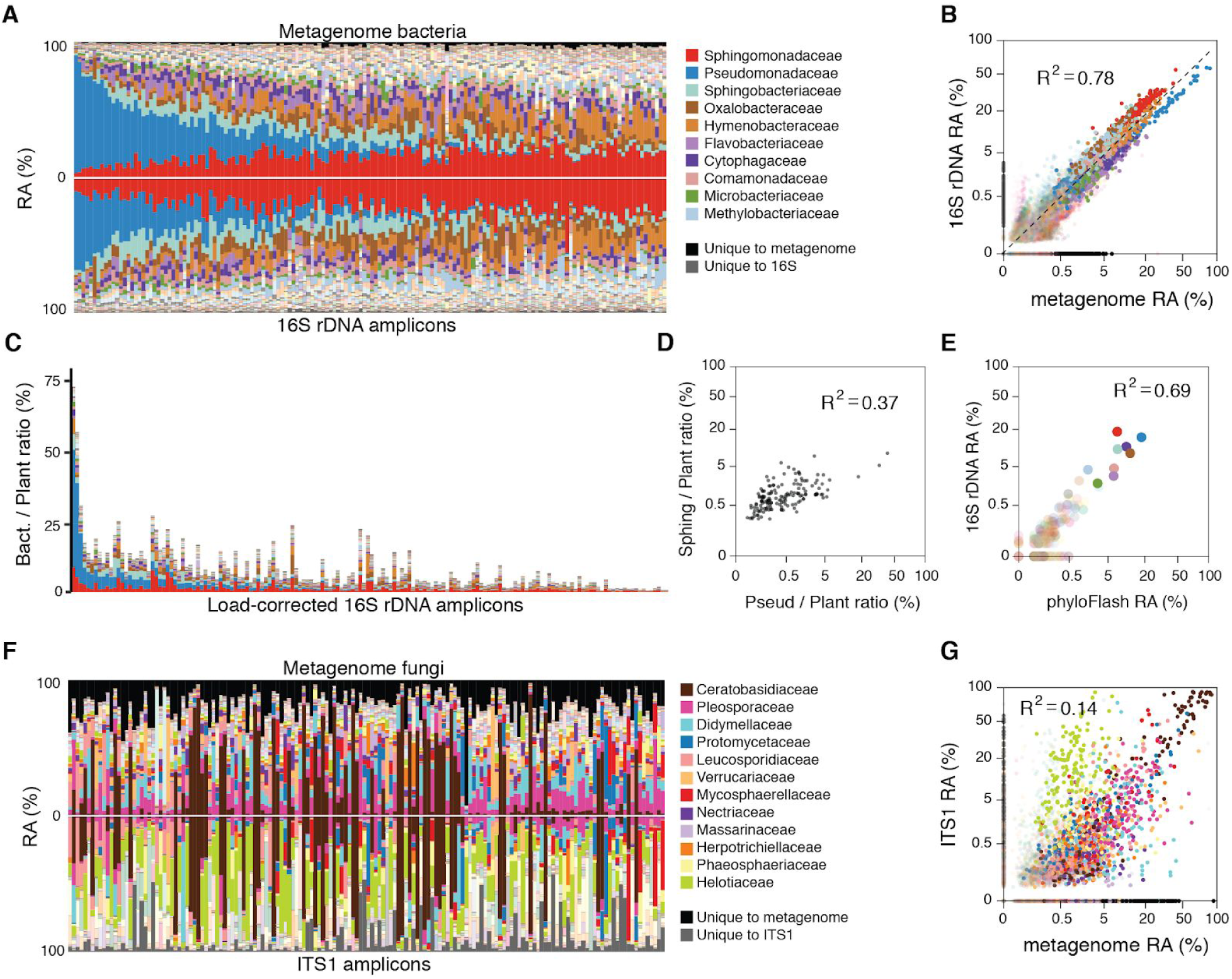
Enhancing utility of metagenome data with parallel amplicon data. **(A)** The relative abundances (RA) of bacterial families as determined by the shotgun metagenome pipeline (top) mirrored against bacterial families as determined by the 16S V4 rDNA amplicon pipeline (bottom) for batch 3 plants (columns). Samples in all panels are ordered by the abundance of Pseudomonadaceae (blue) in the metagenome. Taxa unique to the metagenome are shown in black, those unique to amplicons are shown in dark gray. **(B)** The two data sets from panel A, fourth root transformed and shown as a scatterplot. The dotted line represents perfect correlation. **(C)** The amplicon data from panel A, bottom, scaled by common taxa shared between the metagenome and amplicon data. **(D)** The data in panel C, fourth root transformed and shown as a scatterplot. **(E)** Scatterplot of fourth root-transformed bacterial family abundances, comparing 16S rDNA amplicon data to 16S rDNA sequences detectable in the metagenome (using phyloFlash). Same color scheme for families as in panels A-D. **(F)** The relative abundances (RA) of fungal families as determined by the shotgun metagenome pipeline (top) mirrored against fungal families as determined by the ITS1 rDNA amplicon pipeline (bottom) for batch 3 plants (columns). Samples are ordered as in panel A. Taxa unique to the metagenome are shown in black, those unique to amplicons are shown in dark gray. **(G)** Scatterplot of fourth root transformed data from panel F.

The concordance between fungal families deduced either from ITS1 amplicons or metagenomes was weaker than for bacterial families (Fig. 4F,G), with a Pearson coefficient of correlation R^2^ of 0.14. Several factors could explain this difference. First, fungi are less abundant overall, meaning their quantification is based on fewer sequences and therefore noisier. In agreement, the Pearson correlation coefficient R^2^ of metagenome versus amplicon data for the most abundant fungal family, Ceratobasidiaceae, was much higher than the average, with R^2^ = 0.88. Among other families, the Helotiaceae were especially poorly correlated, and were far more abundant in the ITS1 data (Fig. 4F,G). While this could be due to a bias of the ITS1 primers for this family at the exclusion of others, it could also be that metagenome sequences from this family are more often erroneously assigned to other, sequence related families, deflating counts of Helotiaceae. Another source of noise between the datasets could come from the fact that fungal genomes vary more widely than bacteria in size, and the fact that rDNA copies are not well correlated with fungal genome size [45]. This can introduce biases because organisms with larger genomes would appear to have higher abundances in metagenomes due to more mapped reads. Because these larger genomes may not have more rDNA copies, ITS1 amplicon counts are less affected by differences in genome size. Additionally, because a much smaller fraction of fungal genomes – as compared to bacterial genomes – codes for proteins, and because gene number in fungi varies much less than genome size, many fungal sequences in the metagenome may not be represented in the protein databases used for classification and quantification.

The close concordance between metagenome and 16S rDNA relative abundances enables the scaling of 16S rDNA amplicon data based on bacterial load obtained from metagenome data. Using amplicons, abundant plant host DNA is much more easily blocked using either PNA oligomers [46] or blocking primers [47], meaning that for samples with high plant DNA content, amplicon sequencing can sample many more microbes at a lower cost than metagenome sequencing. Entries in amplicon databases currently represent the taxonomic breadth of microbes more evenly than whole genome or protein databases, and therefore may provide more consistent classification [48]. In addition, we have shown above that relatively shallow metagenome sequencing is sufficient to find enough classifiable reads with which to estimate the bacterial load of a sample (Supplementary Figure 18). Low-cost shotgun library preparation methods [49] in particular make a hybrid approach, in which amplicon sequencing is combined with low-depth shotgun sequencing, attractive. We used the bacterial load as calculated from plant-scaled metagenome data to adjust the abundances of 16S rDNA data to reflect estimated loads (Fig. 3C). As would be expected from the close correlation of 16S rDNA and metagenome data, the adjusted 16S rDNA data accurately captured the slightly positive correlation between Pseudomonadaceae and Sphingomonadaceae across the dataset (Fig. 3D).

## Discussion

Describing the phyllosphere-associated microbial community in the context of natural or field cultivated plant populations is of fundamental importance for understanding and designing microbial interventions in conservation and agriculture. For years, as in studies of other microbial communities, this has been approached via isolation and culture of specific leaf microbes [50–52]. With the advent of high-throughput amplicon and short read sequencing, it has become easier to address the larger community of taxa that interacts with its host. Here, we investigated the advantages as well as the limits of whole metagenome shotgun sequencing to study the leaf microbiota.

A first finding was that de novo assembly produced few longer sequences, and thus did not provide advantages over directly mapping short reads to reference databases for taxonomic assignment. Using taxonomic assignments inferred from short reads directly, we found that the relative abundance of bacterial taxa among all bacteria was highly consistent with 16S rDNA amplicon measurements, with the highest correlation seen for the most abundant taxa. The relative abundance of fungi in the metagenome correlated less well with the ITS1 amplicons, which could be explained at least in part by their lower abundance as well as more complex genomes of fungi compared to bacteria. Nevertheless, the metagenome data clearly showed that fungi are ubiquitous in *A. thaliana* leaves, even though they are usually only a minor part of the overall *A. thaliana* phyllosphere microbiome.

We used shotgun sequencing to estimate microbial load across samples, which varied substantially. The absolute estimates allowed us to reveal the extent to which normalizing bacteria or fungi to a common value via rarefaction or by total sum scaling, as is common practice, may mislead researchers to equate an increase or decrease in relative abundances with a change in absolute abundances.

We used load corrected bacterial taxonomic profiles to explore similarities and differences among the microbiomes. We did not detect a strong individual influence of either environment or host genetics on the structure of leaf communities, although some combination of both contributed. We found that the most abundant taxa in a sample predicted the community structure of the other microbes in the sample. In our natural host populations, there was not enough genetic diversity at each site to allow us to disentangle the effects of site and host genotype on microbiome composition. Whether variation between genetically identical hosts reflects only stochastic effects, or variation in microenvironment, needs to be determined.

An important goal of microbial ecology is to uncover specific interactions between community members, which can point to key taxa that have a major effect on community composition [53,54]. Correlation networks present a valuable tool for such investigations; we have demonstrated that correlations based on relative abundances can lead to very different networks than correlations based on absolute abundances. In our specific case, relative abundance data had suggested that Pseudomonadaceae and Sphingomonadaceae, two of the most common bacterial families typically found in a leaf microbiome, were negatively correlated, when in fact they were positively correlated, as demonstrated with the absolute abundance data. We observed such patterns for several other taxa pairs as well. It is striking that statistically significant correlations were always positive in our dataset when using load corrected data. This could simply reflect generally stable relationships between community members, such that they tended to succeed or fail together as they colonized the plant.

In conclusion, we have demonstrated the advantages of using metagenome shotgun sequencing either alone or in combination with 16S rDNA and ITS1 amplicon sequencing for measuring microbial communities in *A. thaliana* leaves. Modest read depth, as few as 300,000 reads per sample, is sufficient to enable quantitative taxonomic assignment that is comparable to amplicon sequencing. In addition, it turns out once more that the small genome of *A. thaliana* is a substantial advantage, as it may currently be cost prohibitive to extend our direct metagenomic approach to other species. This is yet another reason to use *A. thaliana* (or other species with relatively small genomes) for microbiome studies in ecological settings. On the other hand, the hybrid approach of using lower coverage metagenome data to estimate microbial load and to use this information to scale amplicon data may cost effectively support endophytic analysis of plants with larger genomes as well.

## Methods

### Sampling, Processing of Plants, Metagenomic Library Preparation

Plants were sampled from previously-described sites Eyach (EY), Pfrondorf (PFN) and Jugendhaus Einsiedel (JUG) around Tübingen, Germany [15,55], in four distinct sampling batches which also had different processing details, representing our evolving pipeline.

#### Batch 0 – Single plant

A plant visibly infected with both *Hyaloperonospora arabidopsidis* and *Albugo* sp. was collected in Fall 2014 from Gniebel (48° 34′ 34.10″ North Lat., 9° 10′ 55.42″ East Long.) using sterile tweezers and scissors, placed in a sterile 15 mL tube, and brought back to the lab on ice where it was frozen at −80°C until further processing. The frozen plant was ground in the presence of liquid nitrogen using a mortar and pestle that was lined with 4 layers of autoclaved aluminum foil. Approximately 250 g of the resulting powder was used for DNA extraction, using a custom protocol we previously described [55]. Briefly, the sample was subjected to bead-beating in the presence of 1.5% sodium dodecyl sulfate (SDS) and 1 mm garnet rocks, followed by SDS cleanup with ⅓ volume 5 M potassium acetate, and then SPRI beads. The library was prepared using the TruSeq Nano kit (Illumina), with DNA shearing performed with a S2 focused ultrasonicator (Covaris) as suggested in the manufacturer’s protocol. Rather than Illumina adapters, we used custom adapters described in [56]. The sample was sequenced on one lane of a HiSeq 2000 instrument (Illumina), using a 100 bp single-end kit.

#### Batch 1 – Nine plant test of shearing methods

Nine plants were sampled from Eyach in late December 2014. Rosettes were collected in 50 mL tubes with flame-sterilized scissors and tweezers and brought back to the lab for processing. In the lab, 3 rosettes were left unwashed, 3 were washed in sterile water, and 3 were washed in Silwet L-77 solution. Rosettes were then snap frozen and ground to a fine sand-like consistency with sterile aluminum foil-lined mortar and pestles, as described above. For large rosettes, the ground plant material was transferred among up to 3 DNA-extraction tubes which were processed in parallel to better represent the sample, and pooled again prior to library preparation. The DNA was extracted as described above for the Batch 0 plants. Two sets of libraries were made for the nine plants using homebuilt protocols: one sheared via Covaris and one sheared via Shearase enzyme.

##### Covaris based

For one set of libraries, 100 ng of DNA in 130 μL of elution buffer was sheared on a S2 focused ultrasonicator (Covaris) for 65 seconds using intensity = 4, Duty cycle = 10%, and 200 cycles per burst, to yield a fragment size of approximately 350 bp. The sheared DNA was cleaned with SPRI beads in a 0.8:1 bead to sample ratio, and eluted in 15 μL. End-repair, A-tailing, and adapter ligation were performed similar to [57] following “Alternative Protocol 2” with double DNA size selection after the End-repair step, and using homemade SPRI beads instead of AMPure XP beads. Other minor modifications were that the total volume of the end repair reaction was scaled down to ¼ volume, with DNA eluted after SPRI-cleanup in 17 μL. The total volume of the A-tailing reaction was scaled down to ½ volume. Again, custom adapters described in [56] were substituted for Illumina adapters.

##### Shearase based

For the second set of libraries, 100 ng of DNA in 20 μL of EB buffer was mixed with 9.5 μL of 3X reaction buffer and 0.5 μL of dsDNA Shearase Plus, and incubated for 30 min at 37°C to yield a size range between 200-1000 bp, before the reaction was stopped by addition of 3 μL EDTA. The shared DNA was cleaned with SPRI beads and eluted in 17 μL EB. Size selection, A-tailing, and adapter ligation were performed exactly as with the Covaris based protocol. Final cleaned libraries prepared using both Covaris and Shearase protocols were quantified with PicoGreen (invitrogen) using 1 μL of DNA in 100 μL reactions, and molecules were pooled in equimolar amounts. The pooled library was size selected for fragments between 350 and 700 bp on a Blue Pippin instrument (Sage Science, Beverly, MA, USA). All samples were sequenced on the Illumina HiSeq 3000 with the 2×150 paired end protocol.

#### Batch 2 – Set of 90 plants

Plants were collected from EY and PFN in Fall 2014 (Nov. 24 and 25) and Spring 2015 (March 18 and 19). All samples were brought back to the lab in 50 mL tubes, washed 3x in sterile water to remove adhering dust and soil, and then flash frozen and stored at −80 °C until they were ground in sterile foil-covered mortar and pestle. DNA was extracted as described for other plants above. Metagenomic libraries were prepared as described for Covaris-sheared libraries from Batch 1; the entire set of 90 libraries was quantified, combined to one pool, size selected, and sequenced with 2×150 paired ends over lanes of a HiSeq3000 instrument.

#### Batch 3 – Set of 176 plants

Plants were harvested from EY (11 Dec. 2015 and 23 Mar. 2016), JUG (15 Dec. 2016 and 31 Mar. 2016) and PFN (31 Mar. 2016). Whole rosettes were removed with sterile scissors and tweezers, and washed 3x with sterile water. Two leaves were removed and independently processed to culture bacteria as previously published in [55], and the remaining rosette was flash-frozen on dry ice and processed for metagenomic sequencing and 16S rDNA sequencing of the V4 region. The metagenomic libraries were prepared using a modification of the Nextera protocol for smaller volumes similar to [58], as previously described [55]. As for Batch 1 and Batch 2 plants, the full set of 176 libraries was quantified and combined to one pool, size selected for 350 - 700 bp final library size, and sequenced with 2×150 paired ends over multiple lanes of a HiSeq3000 instrument.

### 16S rDNA V4 library construction and sequencing for Batch 3 plants

Amplicon sequencing of the V4 region of the 16S rDNA gene performed using a two step PCR protocol using PNAs to block chloroplast and mitochondrial sequences, slightly modified from [46]. The first PCR step amplified the rDNA using 515F [59] and 806R [60] primers as well as short overhangs (Supplementary Table 1) and a second step primed these overhangs to added Illumina adapters. The primers differed from Lundberg et al. 2013 in two key ways. First, although the frameshifting nucleotides were kept, the molecular tagging nucleotides were removed from the primers to make the protocol simpler and robust to more variable DNA quantities. Second, the primers were modified such that the Illumina TruSeq priming sequences were used on both the forward and reverse primers, as opposed to the use of a Nextera sequence on the forward primer. Unique barcoding of samples was accomplished by use of 96 independent indexing primers in the second PCR, combined with two combinations of frameshift primers in the first PCR as explained in [46]. Half of the samples from the first PCR were amplified with 515F frameshifts 1, 3, and 5 paired with 806R frameshifts 2, 4, and 6. The other half of the samples from the first PCR paired 515F frameshits 2, 4, and 6 with 806R reverse frameshifts 1, 3, and 5. This strategy allowed up to 192 samples to be uniquely indexed.

In the first PCR, each reaction was prepared in 60 μL, which was split into three 20 μL reactions run in parallel for 29 cycles. Three parallel reactions helps mute the influence of stochastic bias that might affect any single reaction. The 60 μL mix contained 6 μL of TAQ buffer (NEB), 3 μL of 5 μM forward primers mix, 3 μL of 5 μM reverse primers mix, 0.45 μL of 100 μM pPNA, 0.45 μL of 100 μM mPNA, 1.2 μL of dNTPs (10 mM), 0.48 μL of Taq polymerase (NEB), 40.4 μL of PCR-grade water, and 5 μL of template DNA. The first PCR was run for 94°C for 2 minutes followed by 29 cycles of 94°C for 30 s, 78°C for 5 s, 50°C for 30 s, and 72°C for 1 min, and finally 72° for 2 min. The three 20 μL reactions were pooled and 5 μL was run on a gel to confirm amplification. The remaining 55 μL were cleaned with 55 μL of SPRI beads [61] at a bead:sample ratio of 1:1 to remove PCR primers the DNA was resuspended in 30 μL of water. Between 1 and 5 μL of this product from the first PCR, based on gel band intensity, was used in the second PCR of 6 cycles to add illumina adapters.

The second PCR was prepared in 25 μL, and contained 5 μL of Q5 PCR buffer (NEB), 0.0625 μL of 100 μM universal forward primer, 1.25 μL of 5 μM barcoded reverse primer, 0.5 μL of 10 mM dNTPs, 0.25 μL of Q5 polymerase (NEB), 12.875 μL of PCR-grade water, and 5 μL of water + DNA from the first PCR. The second PCR was run for 94°C for 1 minute followed by 6 cycles of 94°C for 20 s, 60°C for 30 s, and 72°C for 30 s, and finally 72°C for 2 minutes. Successful addition of adapters was confirmed by 5 μL of the final product from each reaction on an agarose gel, allowing visualization of a size shift. Final amplicons averaged 430 bp in length. The remaining 20 μL of product was cleaned with SPRI beads and resuspended in 40 μL of EB. Libraries were quantified by PicoGreen in 100 μL reactions, pooled in equimolar amounts, and sequenced using a MiSeq V2 2×500 reagent kit (Illumina) which was sufficient to overlap and assemble the forward and reverse reads. The frameshifts built into the primers used in the first PCR made the addition of PhiX to increase sequence diversity unnecessary [46].

### ITS1 library construction and sequencing for Batch 3 plants

ITS1 rDNA amplicons were prepared similarly to 16S rDNA amplicons, using gene-specific primers for the first PCR and adding indexes and adapters in the second PCR. We used a protocol modified from [47], which uses blocking primers to prevent amplification of plant sequences. Because blocking primers, unlike PNAs, result in a quantifiable PCR product, we used the cycling conditions suggested in [47] to prevent the product of the blocking product from reaching detectable levels. As with the 16S rDNA protocol, we used six frameshifting forward ITS1F primers and six frameshifting reverse ITS2R primers. The first 60 μL PCR reaction (also run as three parallel 20 μL reactions) included 6 μL of 10X ThermoPol Taq buffer (NEB), 0.96 μL of 5 μM forward primer (0.08 μM final), 0.96 μL of 5 μM reverse primer (0.08 μM final), 0.15 μL of 100 μM forward blocking primer (0.25 μM final), 0.15 μL of 100 μM reverse blocking primer (0.25 μM final), 1.2 μL of 10 mM dNTPs, 0.48 μL of Taq DNA polymerase (NEB), 45.1 μL of PCR-grade water, and 5 μL of DNA. The first PCR was run for 94°C for 2 min. followed by 10 cycles of 94°C for 30 s, 55°C for 30 s, and 72°C for 30 s, and finally 72° for 3 min. The three 20 μL reactions were pooled, cleaned with a bead:sample ratio of 1:1 to remove PCR primers, and resuspended in 30 μL of water.

The second PCR was prepared in 25 μL, and contained 5 μL of Q5 PCR buffer (NEB), 0.0625 μL of 100 μM universal forward primer, 1.25 μL of 5 μM barcoded reverse primer, 0.5 μL of 10 mM dNTPs, 0.25 μL of Q5 polymerase (NEB), 4.875 μL of PCR-grade water, and 13 μL of DNA from the first PCR. The second PCR was run for 94° for 1 minute followed by 25 cycles of 94° for 20 s, 60° for 30 s, and 72° for 30 s, and finally 72° for 2 minutes and cool down to room temperature. Successful PCR and addition of adapters was confirmed by 5 μL of the final product from each reaction on an agarose gel, with the major band produced around 400 bp in length. The remaining 20 μL of product was cleaned with SPRI beads and resuspended in 40 μL of elution buffer. Libraries were quantified with PicoGreen (ThermoFisher Scientific) in 100 μL reactions, pooled in equimolar amounts, and sequenced using a MiSeq V3 2×600 reagent kit (Illumina).

### Amplicon quality processing, clustering, and classification

Raw sequences from both 16S and ITS1 rDNA amplicons were first demultiplexed according to their 9 bp barcodes added in the second PCR, not allowing any mismatches. All sequences were further demultiplexed by the frameshift combinations using strict regular expressions without mismatches in any part of the primer sequence (https://github.com/derekLS1/Metagenome2019). Forward frameshifts 1, 3, and 5 were only allowed pairings with reverse frameshifts 2, 4, or 6. Forward frameshifts 2, 4, and 6 were only allowed pairings with reverse frameshifts 1, 3, and 5.

Forward and reverse reads from the 16S rDNA sequences were merged with FLASH [62] using a minimum overlap set to 30 bp and (-m 30). Most ITS1 amplicons were small enough to overlap with these longer reads, but some reads were longer and overlap was not possible, so only the forward read was used for downstream analyses (read 1), although the frameshift in read2 was used for demultiplexing.

All primer sequences were removed. Because of the frameshifts in the primer sequences, ITS1 read 1 sequences had variable lengths after removing primers, and therefore all were trimmed to a common length of 271 bases before clustering. Additional quality filtering, removal of chimeric sequences, OTU preparation and OTU tables, and taxonomic assignment were done with USEARCH10 [63] (https://github.com/DerekLS1/Metagenome). OTUs were prepared at 100% as ‘zero-radius OTUS’ (zOTUS, a form of Amplicon Sequence Variant) [64]. The 16S rDNA taxonomy was based on the RDP training set v16 (13k seqs.), and ITS1 taxonomy was based on UNITE USEARCH/UTAX release v7.2 (UNITE Community. https://doi.org/10.15156/BIO/587476).

### Metagenome read QC and host data removal

Sequencing libraries were subject to adapter trimming and quality control with Skewer [65]. Reads were trimmed to a minimum length of 30 bp and minimum average Phred score of 20. After sequencing, samples were composed of a mixture of mostly host *Arabidopsis thaliana* reads and microbial origin reads. In order to remove most of the plant reads, libraries were aligned against the *A. thaliana* reference genome [16] using the *bwa mem* algorithm with standard parameters [66]. After mapping, only read pairs for which neither of the mates mapped against the plant reference genome were mapped against the metagenomic reference. Data aligned to the host was later used for host plant genotyping.

### Metagenomic profiling

Using DIAMOND [18] with default mapping parameters, the putative microbial reads were mapped against the entire NCBI nr protein database (March 2018), which includes protein sequences from all three domains of life and viruses. In order to keep analysis time and file sizes manageable, a maximum of 25 matches per sequencing read was permitted. DAA (Diamond analysis archive) files were then parsed for taxonomic binning with MEGAN [20]. Reads were binned to different taxa using the weighted LCA algorithm [19] using only hits that were within 10% of the highest matching score. In summary, of the maximum 25 matches any read could have, only matches whose score was within 10% of the highest score were used for taxonomic placement. Final count tables for different taxonomic levels were produced based on the binning strategy just described.

The taxa counts were then normalized to adjust for sampling depth by first dividing the abundances in each sample by the number of reads mapping to the plant chromosome in that sample. Then, all values in all samples were multiplied by the mean number of chromosomal plant counts across all samples. This can also be represented by the following formula:

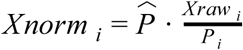

For sample *i*, where *Xnorm* stands for the normalized vector of counts in the sample, 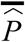 is the mean number of chromosomal plant counts across all samples, *Xraw*_*i*_ is the raw microbial count vector in a sample, and *P*_*i*_ is the number of plant chromosomal reads in that sample.

### Taxonomic correlations and network computation

After computing intermicrobial linear correlations and filtering out weakly associated taxa pairs, network representation was computed with the *networkx* Python package using the *kamada kawai* layout function with standard parameters based on the correlation values.

### Plant genotyping

Individual host genotypes were determined using plant associated reads from each metagenome. Reads aligned to the TAIR10 reference genome [16] were filtered to a minimum mapping quality of 20, resulting in an average genome coverage from 15x to 40x. Single nucleotide polymorphisms were called using FreeBayes [34], and resulting VCF files were filtered using custom scripts. SNPs with a minimum alternative count above 3, minimum read depth of 6, and no more than 5% missing data across all samples were kept for downstream analysis.

For determining genotype groups, a genetic distance matrix was computed with ngsDist [67] from the alternative allele count matrix of all SNPs that passed filtering thresholds. This distance matrix was used as input in *tsne* [68] to visualize sample clustering.

### Adjusting 16S rDNA amplicon data by bacterial load

To adjust the 16S rDNA amplicon dataset to correct for bacterial load, the abundance of each OTU from a sample in the total sum-scaled OTU table can be multiplied by a load scaling factor calculated from the metagenome data for that sample. The simplest load scaling factor is the ratio of all bacteria to plant chromosomal reads in the metagenome sample. If read depth in the metagenome allows, a more precise scaling factor can be calculated based on bacterial families detectable by both methods. Both methods yield similar results in our dataset, because the majority of sequencing reads in both methods fall into bacterial families shared by both methods. We scaled based on bacterial families shared by both methods.

To correct the 16S rDNA dataset by shared taxa in the scaled metagenome dataset, the 16S rDNA dataset was first normalized to 100% in each sample by total sum scaling. The common bacterial families that could be identified by at least a single read in both the metagenomic and 16S rDNA datasets were then identified for each sample, and the sum of read counts in falling in these common taxa was calculated for each sample in both the metagenome and 16S rDNA datasets. The sum of common taxa for each sample in the plant-chromosome scaled metagenome dataset was divided the sum of reads in these common taxa in the corresponding 16S rDNA table to yield a load scaling ratio. The load scaling factor was multiplied by all the 16S rDNA counts in that sample to produce load-corrected 16S rDNA abundances, closely matching the values obtained from the metagenome. For each sample i,

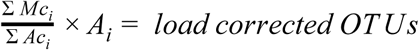

Where Σ*Mc*_*i*_ is the sum of metagenome reads in common taxa from the plant chromosome-scaled metagenome table, Σ *Ac*_*i*_ is the sum of 16S rDNA reads falling in common taxa, and *A*_*i*_ is the full set of 16S rDNA read counts for that sample.

### Comparison of 16S rDNA reads from the metagenome to 16S rDNA V4 amplicons

Metagenome reads from each sample in the Batch 3 dataset were mapped to the RDP 16S rDNA training set using phyloFlash [44]. Each metagenome sample yielded an average of 280 mapped 16S rDNA reads, with many yielding fewer than 100 reads. Because for most samples there were too few reads to compare directly to their 16S rDNA amplicon counterparts, samples were pooled to make one aggregate metagenome dataset containing 52,589 16S rDNA phyloFlash sequences. This was then compared to a corresponding 16S rDNA amplicon dataset comprising 52,589 sequences subsampled from the full 16S rDNA dataset. Each sample contributed a matching number of phyloFlash 16S rDNA or amplicon 16S rDNA reads to either the phylFlash 16S rDNA pool or the amplicon 16S rDNA pool respectively. The family level relative abundances for these pools were then plotted against each other.

## Data availability

All data in this manuscript has been deposited in the European Nucleotide Archive (ENA). It can be accessed under the project number PRJEB31530. At https://www.ebi.ac.uk/ena

## Author Contributions

JR, DL, and DW planned the study and wrote the manuscript. JR, DL, and OD analyzed the data. DL, SK, and KP collected samples and prepared libraries. TK and GS collected samples and commented on the manuscript

## Acknowledgements

We thank Daniel Huson for his support with software implementations and data analysis, Clemens Weiß for his input and suggestions regarding data analysis, and Dino Jolic for helpful ideas and discussions. Supported by a Human Frontiers Science Program (HFSP) Long-Term Fellowship (LT000565/2015-L to DL), ERC Advanced Grant IMMUNEMESIS (340602), the DFG through Collaborative Research Center CRC1101, and the Max Planck Society (DW).

