## Supplementary_Information.PDF for "Combining whole genome shotgun sequencing and rDNA amplicon analyses to improve detection of microbe-microbe interaction networks in plant leaves"

### Table of contents S1

Supplementary Discussion 1: Subtractive hybridization S2
Supplementary Figure 1: Normalization strategies. S4
Supplementary Figure 2: Unmapped reads. S5
Supplementary Figure 3: Individual metagenome assembly statistics. S7
Supplementary Figure 4: Statistics for assemblies of individual and pooled samples. S8
Supplementary Discussion 2: Metagenomic assembly S9
Supplementary Figure 5: Pseudomonas core genome mapping. S10
Supplementary Figure 6: Technical replicates. S11
Supplementary Figure 7: Mapping rate to NCBI nr by sample. S12
Supplementary Figure 8: Kingdom level taxonomic composition. S13
Supplementary Figure 9: Distributions of Shannon diversity. S14
Supplementary Figure 10: Taxonomic composition by site. S15
Supplementary Figure 11: Top bacterial families abundance distributions. S16
Supplementary Figure 12: High-depth plant metagenome. S17
Supplementary Figure 13: PCA ordination by sampling season. S18
Supplementary Figure 14: PCA ordination by abundant taxa. S19
Supplementary Figure 15: PCoA ordination of MASH distances. S20
Supplementary Figure 16: Correlation matrix of abundant taxa. S21
Supplementary Figure 17: Taxa correlations for bacterial load. S22
Supplementary Figure 18. Depth of sequencing to accurately determine microbial load. S23
Supplementary Figure 19: PCA ordination of fungal family abundance. S24
Supplementary References S25

### Supplementary Discussion 1: Subtractive hybridization

Subtractive hybridization was attempted as a method to enrich finished Illumina libraries for microbial DNA. It is the reverse of hybridization capture, a common and successful enrichment technique to enrich a library for fragments which are similar to the known sequences used in the bait [1]. Unlike hybridization capture, where one keeps that DNA which binds to the bait, in subtractive hybridization one is interested in what does not bind to the bait.

*A. thaliana* seeds were surface sterilized and used to grow microbe-free *A. thaliana* seedlings on agar plates. Week old seedlings were harvested under sterile conditions and used to make microbe-free DNA using sterile reagents. The pure plant DNA was then biotinylated, denatured, and then allowed to hybridize with test DNA library that was made from a mix of both E.coli DNA and sterile-grown *A. thaliana* DNA for a defined period. Different ratios of bait to prey library were used. Finally, the before the biotinylated DNA was removed from the solution using an excess of streptavidin-coated magnetic beads in an attempt to "fish" out the hybridizing plant sequences, leaving a microbe-enriched fraction. In most cases, we used 500 ng of biotinylated plant DNA as bait for 20 ng of library, a 25-fold excess of bait, in a hybridization volume of 9 µL. Total DNA quantification of the reaction mixture showed that streptavidin-coated beads removed most of the DNA from the solution, so 1PCR was used to quantify enrichment of microbial sequences. A titration of *E. coli* DNA into sterile plant DNA could be detected quantitatively using 1PCR, meaning the qPCR assay functioned as expected. Although the beads captured DNA that was enriched for plant DNA, the remaining library was not reliably enriched for microbial sequences, with at best 2.5 fold enrichment. After multiple attempts varying capture parameters, there were no improvements.

Acknowledging the possibility that the streptavidin-coated beads could not remove 100% of the bait DNA from the hybridization, and that this bait DNA was contaminating the qPCR, we decided to run some of the putatively microbe-enriched library on an Illumina MiSeq nano flow cell and count the ratio of plant reads to microbial reads. Because the bait DNA did not have sequencing adapters, it would not be sequenced on the MiSeq, even if it remained in the library. The MiSeq results showed no better enrichment than the qPCR.

We originally attempted subtractive hybridization to achieve drastic microbial enrichment. After achieving at best modest enrichment, we decided that tweaking the protocol to achieve consistent modest microbial enrichment would not be worth the effort, because of the added difficulty of a hybridization step before shotgun sequencing and the unknown potential biases that might be introduced, for example from inadvertently hybridizing microbial sequences that had sequence similarity with parts of the plant genome.

This discussion is not meant to show that subtractive hybridization cannot be effective, but rather to explain that it was not effective in our hands after multiple attempts and should not be approached blithely. For someone with greater experience in similar techniques, it may be entirely feasible and we can share more details of the specific methods we tested.

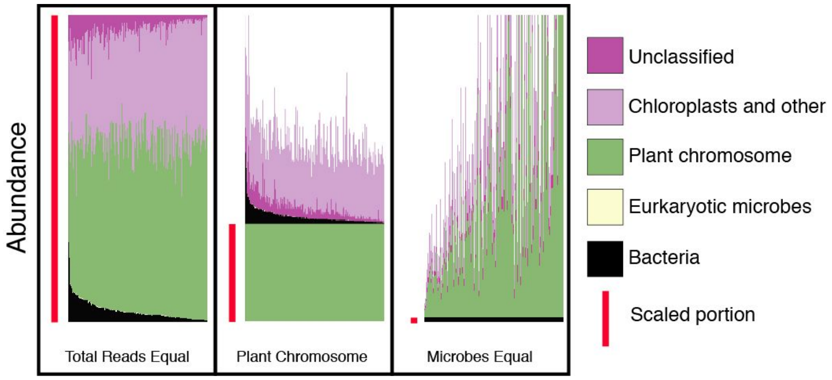


**Supplementary Figure 1: Normalization strategies.**

Shown are three possible normalization strategies to correct for sequencing depth. When total reads are held equal (left), the reads assigned to each taxon are divided by the total number of high quality reads in that sample, such that the sum of all taxa in each sample is 1. When microbes are held equal (middle), the reads assigned to each taxon are divided by the total number of microbial reads in that sample, such that the sum of all microbes in each sample is 1. When plant chromosomal DNA is held equal (right), the reads assigned to each taxon are divided by the number of reads mapping to the plant genome, such that the total number of reads assigned to plant in each sample is 1, and microbial reads across samples may vary.

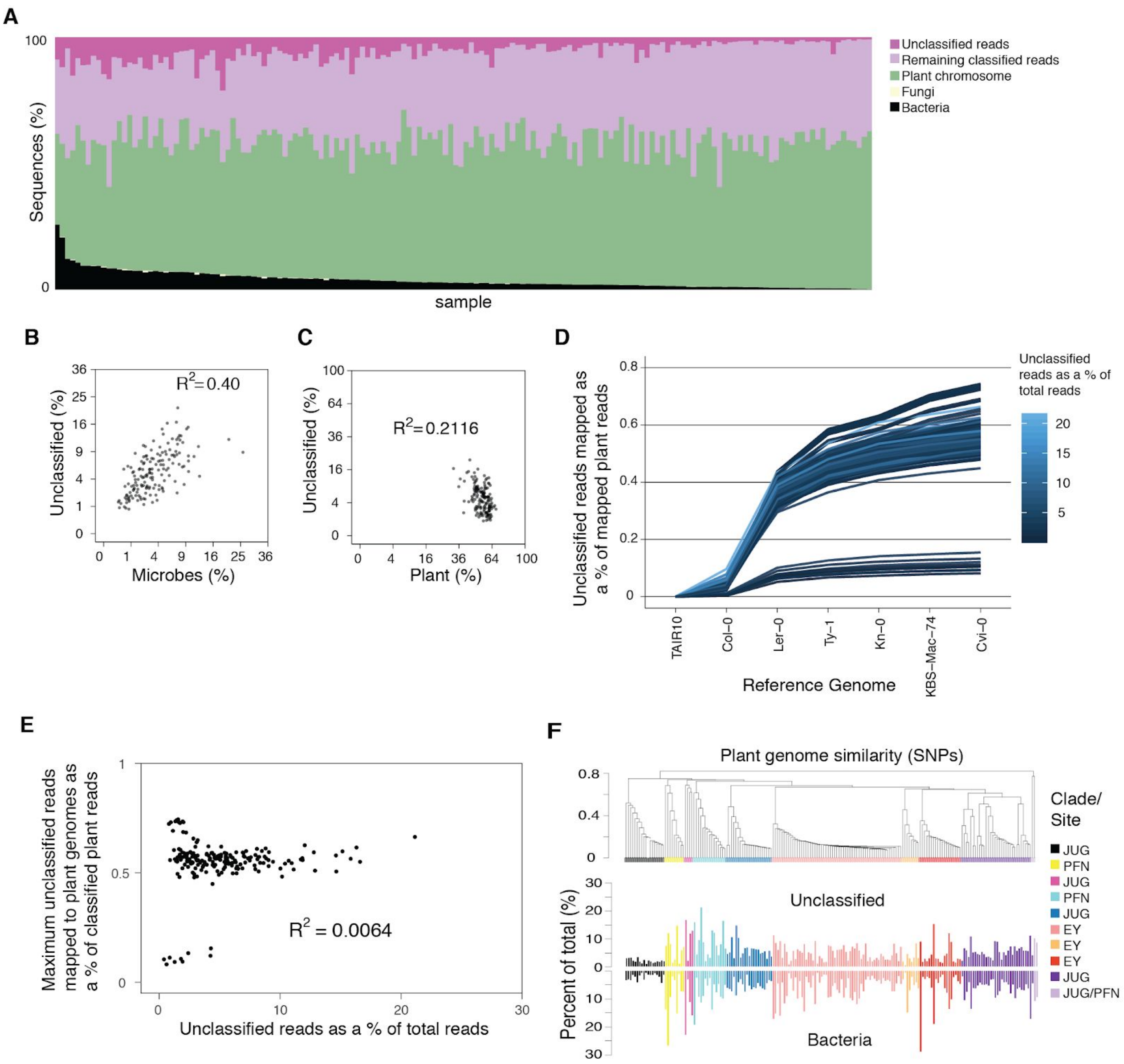


### Supplementary Figure 2: Unmapped reads.

(**A**) Composition of each sample in the batch 3 dataset. Of all reads, the number of bacterial reads (bottom, black) is similar to the number of unmappable reads (top, dark magenta). (**B**) The abundance of unclassifiable reads is positively correlated with microbial (bacterial plus fungal) reads. (**C**) Unclassifiable reads and plant chromosomal reads are weakly negatively correlated. (**D**) Some unclassified reads from each sample in the batch 3 dataset map to nuclear genomes from other *A. thaliana* reference genomes. After mapping sequentially to another Col-0 reference and five additional genomes, the additional mappable reads remained less than 1% of the total classifiable plant reads in each sample. This was true regardless of the relative proportion of unclassifiable reads in the sample, which ranged from less than 0.5% to over 20% (color key). (**E**) The maximum relative read mapping of unclassifiable reads to the additional *A. thaliana* reference genomes (shown above for Cvi-0) in **D** was plotted against the percentage of unclassifiable reads of total reads in each sample. Regardless of the quantity of unclassifiable reads in a sample, the mapping to additional plants was essentially invariant at around 0.5% of total classifiable plant reads for most samples. (**F**) Some closely related plant genotypes, particularly a group from JUG (black), had fewer unclassified reads than others, but the number of unclassified reads was mirrored by a similarly low number of microbial reads.

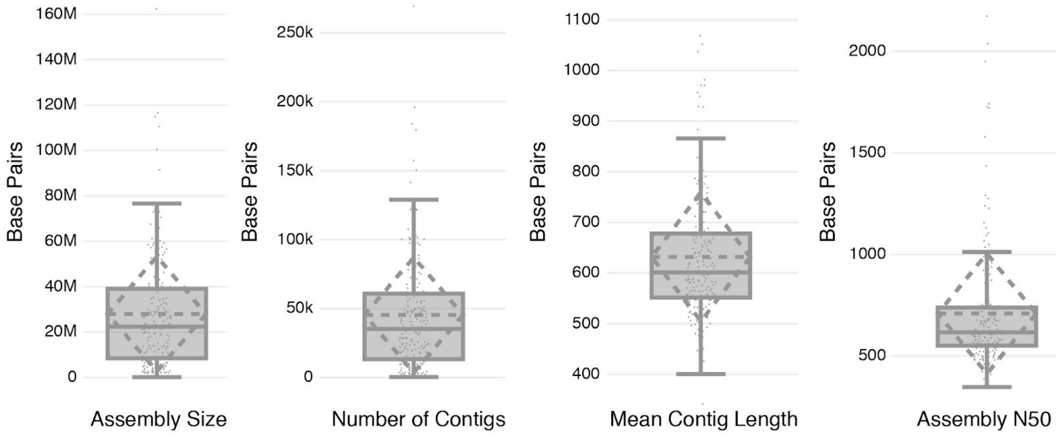


**Supplementary Figure 3: Individual metagenome assembly statistics.**

Distribution of assembly statistics for individual metagenomic assemblies across all datasets. Plant microbiome metagenomes for each sample assembled individually using megahit with "--sensitive" preset parameter and minimum contig length of 200 bp.

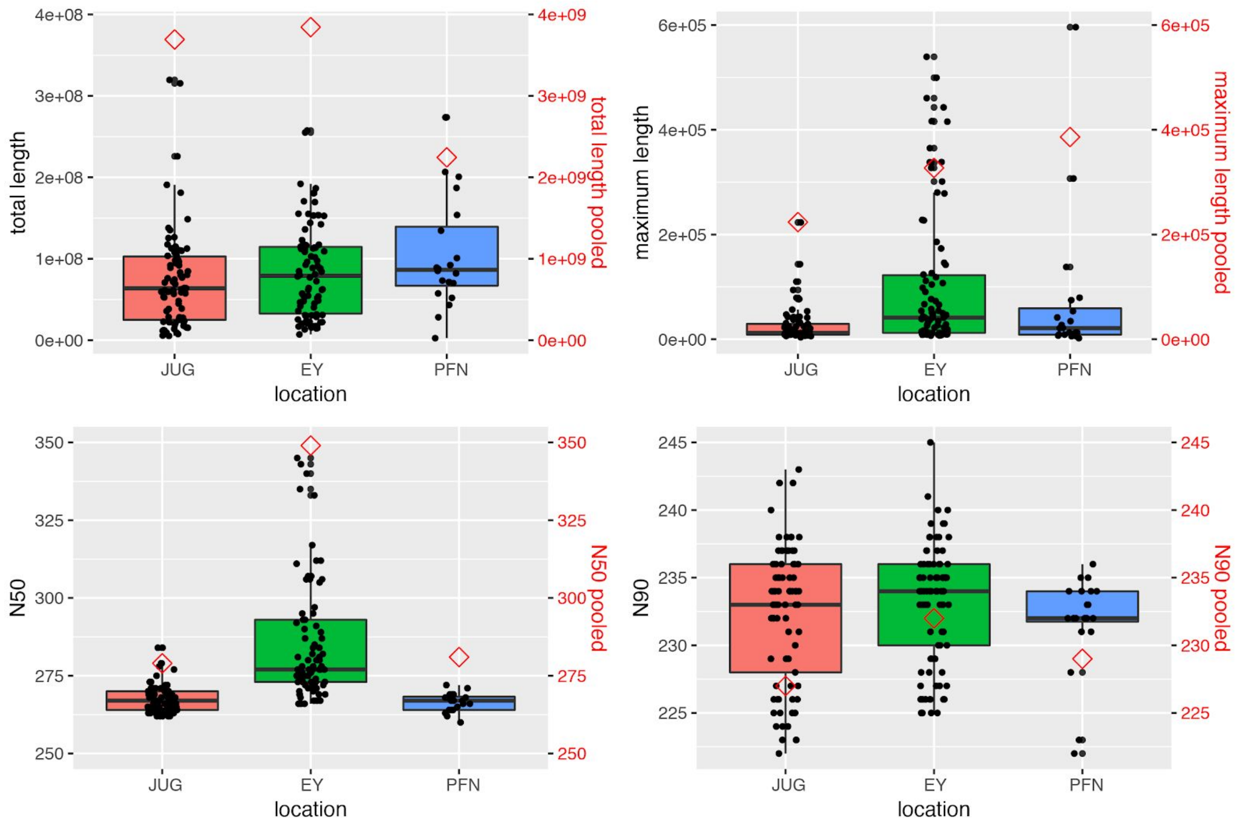


**Supplementary Figure 4: Statistics for assemblies of individual and pooled samples.**

Total length is the sum of all contigs, maximum length is the size of the longest contig, and N50 and N90 give the contigs that make up 50 and 90% of the entire assembly. The box and whisker plots illustrate assembly statistics for 176 individual samples summarized by location. Red diamonds show results for samples pooled by location prior to assembly (different scales used for number of contigs and total length). Abbreviations: Jug - Jugendhaus, Ey - Eyach, Pfn - Pfrondorf.

### Supplementary Discussion 2: Metagenomic assembly

To begin to understand potential reasons for the disappointing success of assembly efforts, we made use of the fact that *A. thaliana* phyllosphere samples in this area of Germany are dominated by a specific clade of closely related *Pseudomonas* strains [2]. We took the 10 samples with the highest number of *Pseudomonas* assigned reads, and we mapped both assembly contigs and raw reads against the core genome of a representative member of this *Pseudomonas* clade. Of the individual reads that mapped to the *Pseudomonas* core genome, around 80% could be classified to the genus *Pseudomonas* using DIAMOND with the NCBI nr database as a reference, vs. 74% of reads associated with contigs mapping to the core genome (Supplementary Figure 5). To reveal the origin of these assembly-associated reads that did not map to *Pseudomonas*, we tested mapping to other microbial clades. There were no core-mapped reads positively classified as belonging to another microbial clade, compared to 0.5% of reads associated with mapped contigs. Around 6% of core-mapped reads were not classifiable, but for reads associated with mapped contigs, this jumped to 10%. These results show that contigs mapping to the core genome include reads from other taxa. These contigs could either be purely other taxa that can also map to a homologous region in the *Pseudomonas* core genome, or they could be chimeric assemblies between Pseudomonas sequences and other taxa; either way, such contigs interfere with further assembly because they can integrate into the assembly and introduce dead ends. Around 6% of core-mapped reads were not classifiable, but for reads associated with mapped contigs, this jumped to 10%, further supporting the idea that unclassified reads are microbial in origin.

Impressive progress has been made with generating assemblies of draft genomes from complex metagenome samples [3], primarily through creative use of binning approaches [4]; a prerequisite for this is, unfortunately, that the primary assembly step generates contigs of sufficient sizes for merging into draft genome assemblies. Due to the complexity of phyllosphere metagenomes, longer sequencing reads are likely necessary to overcome the problem of dead ends introduced by sequence-related regions in other organisms.

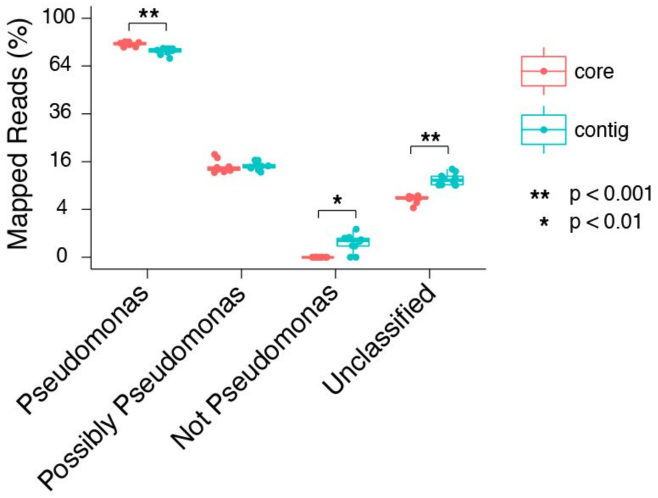


**Supplementary Figure 5: *Pseudomonas* core genome mapping.**

From the 10 samples with the most total *Pseudomonas* reads, single reads were mapped to a *Pseudomonas* core genome and these were classified with DIAMOND (red). Alternatively, reads were assembled into contigs which were in turn mapped to the core genome, and reads associated with these contigs were classified with DIAMOND (blue). Significant differences in the fraction of single reads vs. contig-associated reads that were *Pseudomonas*, consistent with *Pseudomonas*, not *Pseudomonas*, or unclassifiable were determined by a paired Wilcoxon signed-rank test (* = p<0.01, ** = p<0.001).

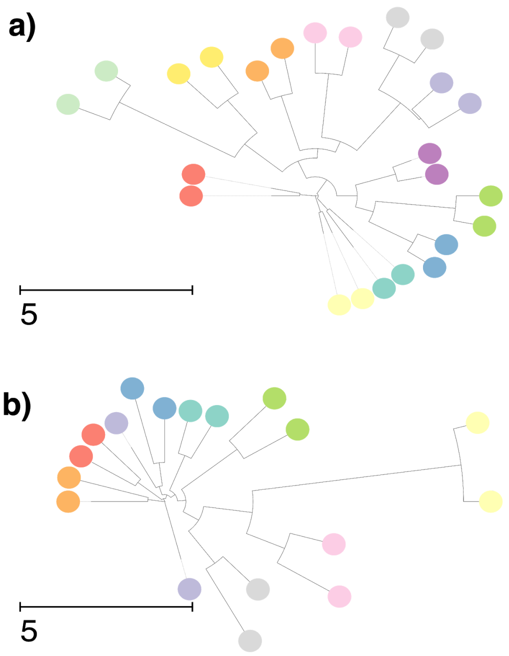


**Supplementary Figure 6: Technical replicates.**

Neighbor joining tree on Euclidean distance matrix of subsampled replicated samples. Microbial fraction of metagenomes was subsampled to 300 000 reads. Taxonomic profiles were recomputed as described in methods and neighbor joining trees were computed based on the Euclidean distance matrix for (**A**) Twelve plants where starting material was divided in two and each half processed separately. (**B**) DNA from batch 1 plants was extracted, and DNA was processed into two sequencing libraries for each plant. Tips of the trees correspond to samples and colors correspond to plant of origin. The plant of origin is recovered for technical replicates.

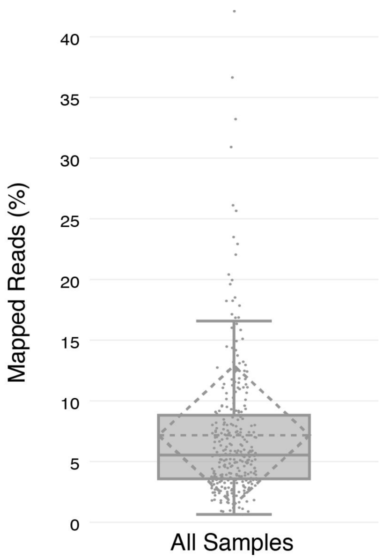


**Supplementary Figure 7: Mapping rate to NCBI nr by sample.**

Distribution of fraction of total reads with high quality mappings to reference database. Solid line indicates median, and dotted lines correspond to mean and standard deviation. Individual data points are plotted for all 275 samples.

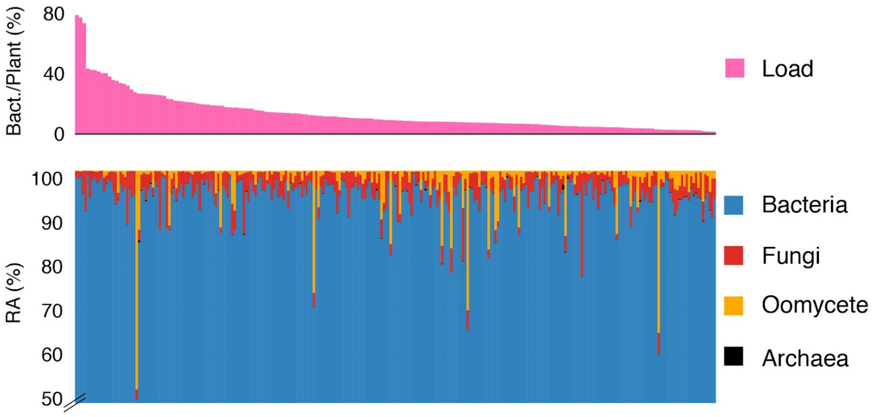


**Supplementary Figure 8: Kingdom level taxonomic composition.**

High level taxonomic composition of leaf microbiomes across all samples. Top panel, microbial load as the ratio of microbial to plant chromosomal reads. Bottom panel, relative abundance of at high taxonomic levels.

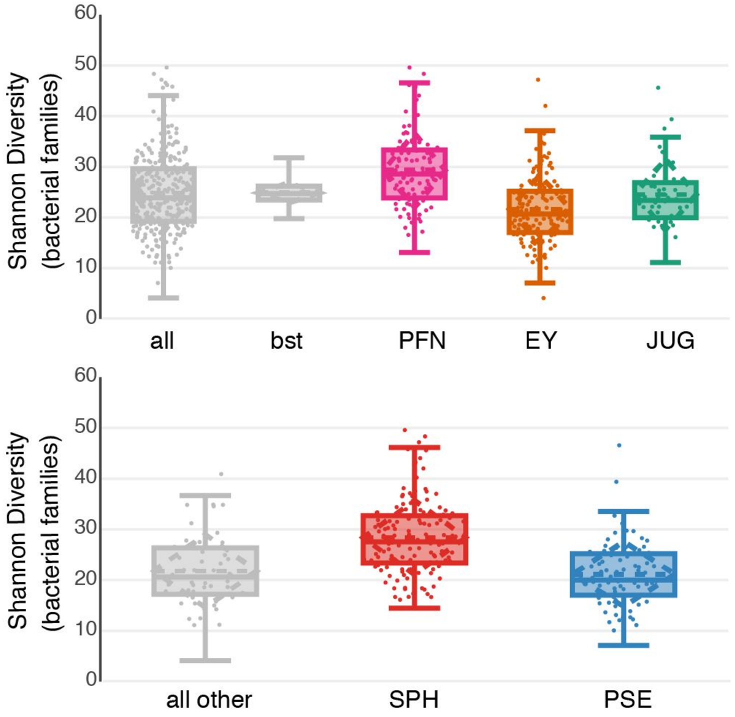


**Supplementary Figure 9: Distributions of Shannon diversity.**

Top, all - entire dataset, bst - 1000 replicates of 15 sample subsampling, PFN: Pfrondorf, EY - Eyach, JUG - Jugendhaus. Bottom, all other - samples with most abundant taxa other than Pseudomonadaceae or Sphingomonadaceae. Sph - Sphingomonadaceae, Pse - Pseudomonadaceae. Shannon diversity (exponential of the Shannon index) in bottom panel computed after removing most abundant taxa.

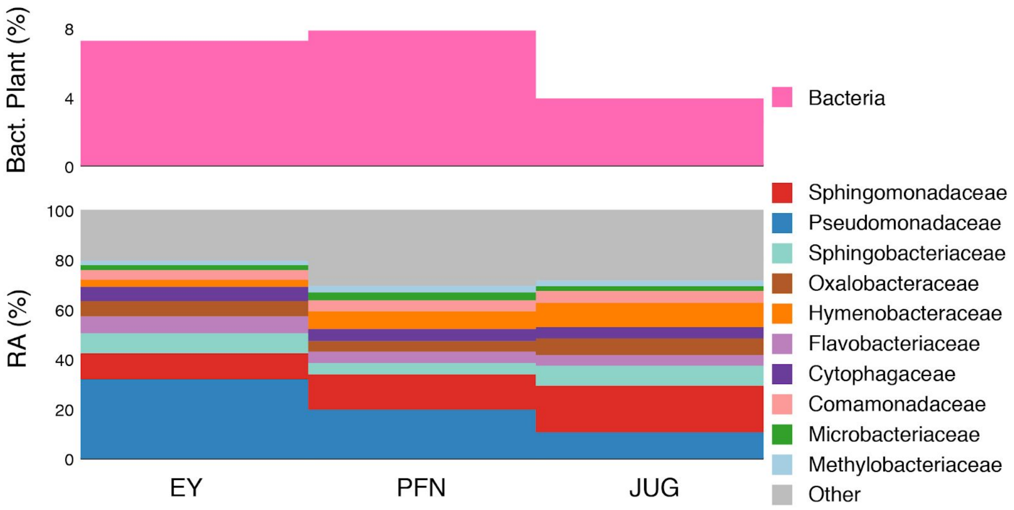


**Supplementary Figure 10: Taxonomic composition by site.**

Top, average bacterial load. Bottom, average relative abundance of bacterial families.

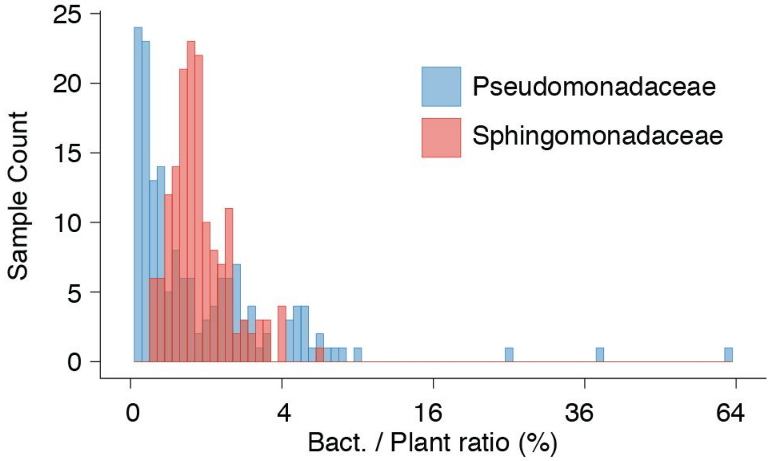


**Supplementary Figure 11: Top bacterial families abundance distributions.**

Pseudomonadaceae have a wider and more extreme distribution than Sphingomonadaceae, with more samples having either extremely low or extremely high Pseudomonadaceae abundances. Based on plant-scaled data.

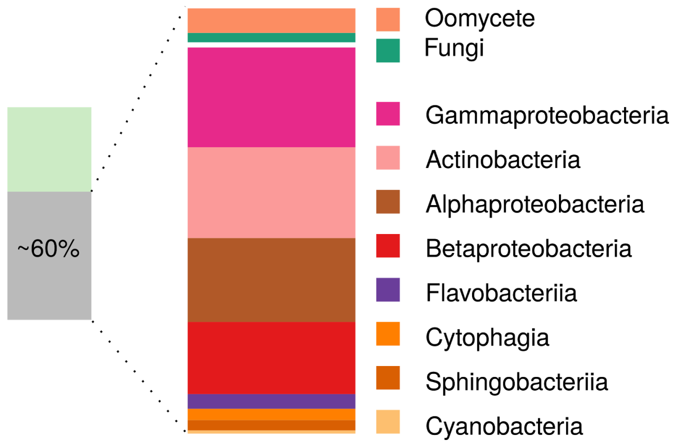


**Supplementary Figure 12: High-depth plant metagenome.**

Taxonomic composition of individual *A. thaliana* leaf microbiomes sequenced at high depth. Left - Fraction of total reads considered non-host. Right - taxonomic decomposition at the family level.

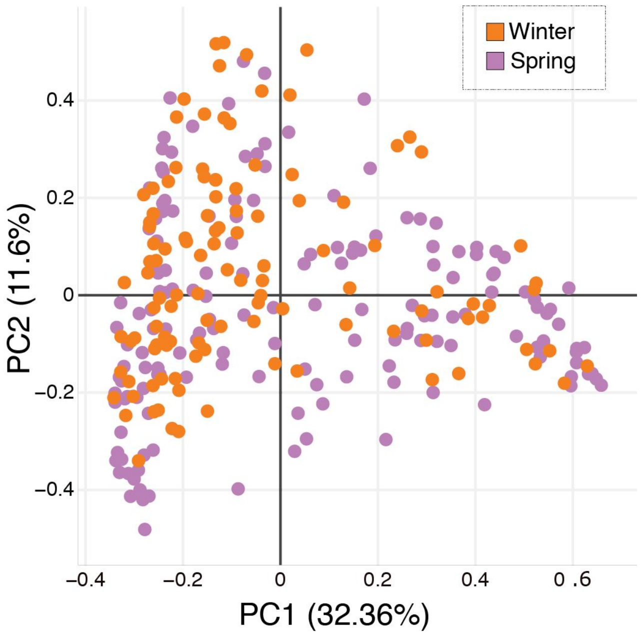


**Supplementary Figure 13: PCA ordination by sampling season.**

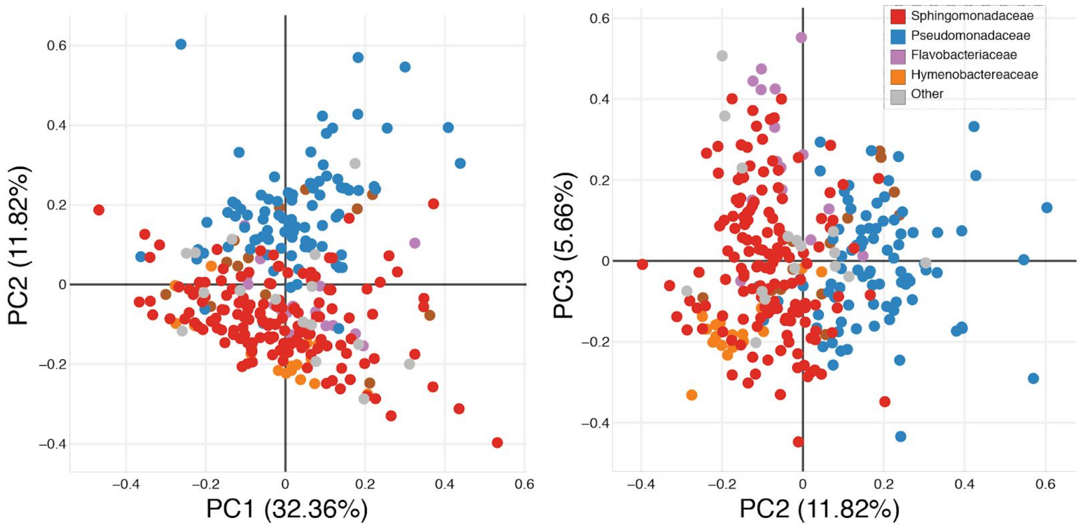


**Supplementary Figure 14: PCA ordination by abundant taxa.**

Family level plant-scaled fourth root transformed count matrix. Samples are colored by most prevalent microbial family.

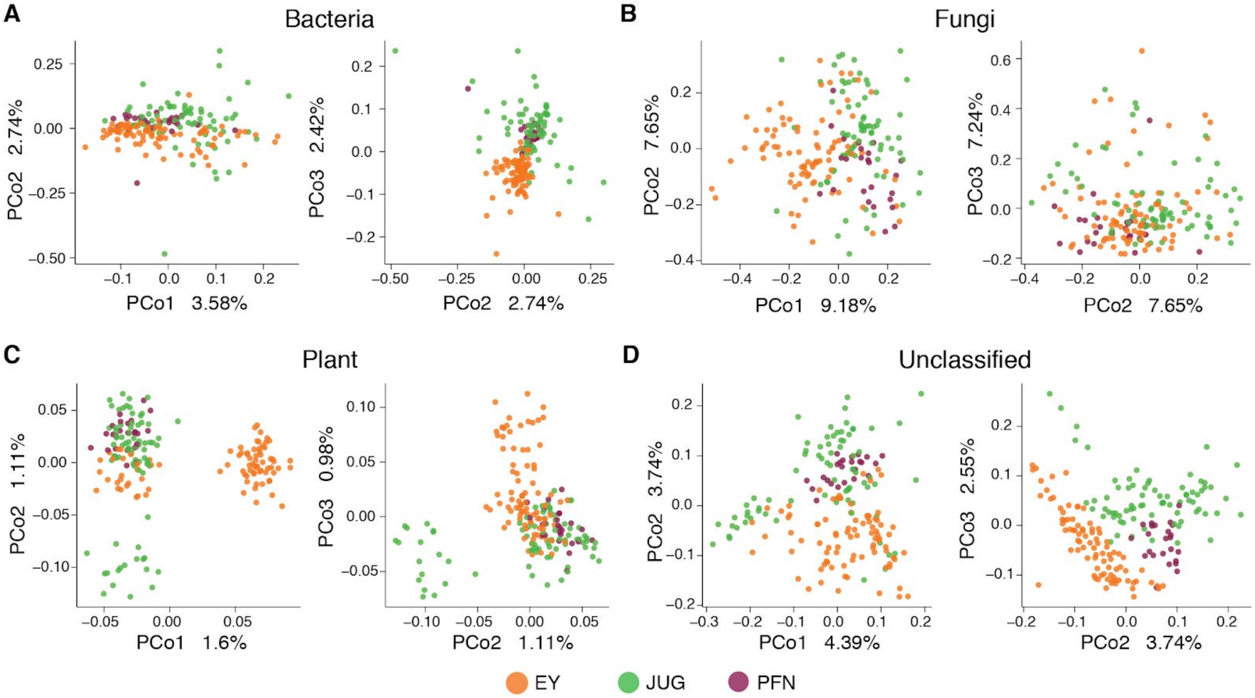


**Supplementary Figure 15: PCoA ordination of MASH distances.**

Reference databases were only used to classify reads as plant, bacteria, fungi, or unclassified. The distance between samples was then calculated by comparing unique k-mers in a reference-independent manner with MASH **[5]**. (**A**) Ordination of square root-transformed MASH distances calculated based on reads classified as bacteria and colored by collection site. (**B**) Same as **A**, but for reads classified as fungi. (**C**) For reads classified as *Arabidopsis thaliana.* (**D**) For unclassified reads.

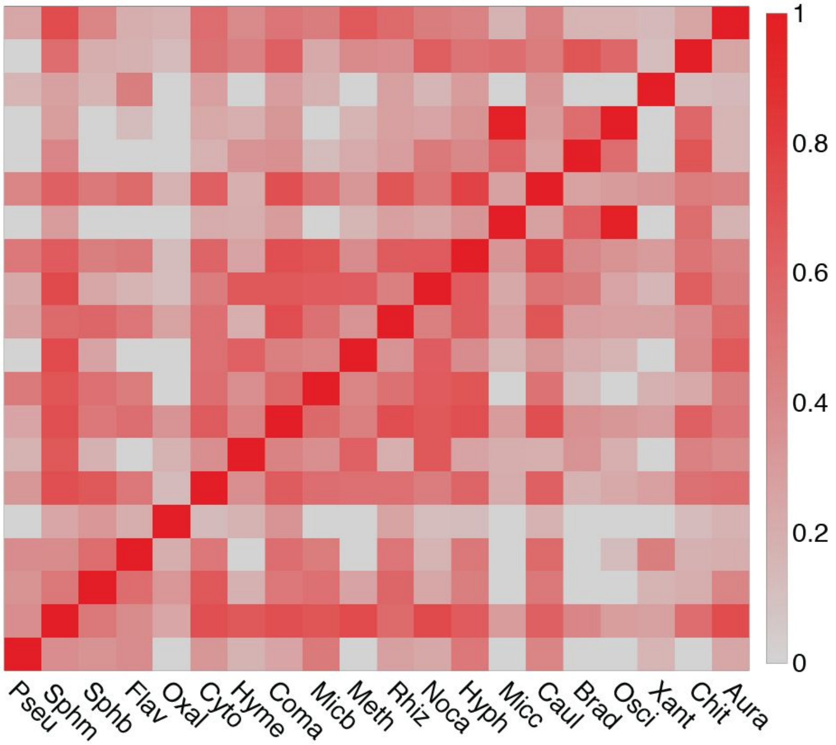


**Supplementary Figure 16: Correlation matrix of abundant taxa.**

Heatmap of taxa correlations measured on read counts. Correlations with p-values below 0.05 after t-test are shown for top 20 most abundant taxa.

Pseu - Pseudomonadaceae, Sphm - Sphingomonadaceae, Sphb - Sphingobacteriaceae. Flav - Flavobacteriaceae, Oxal - Oxalobacteraceae, Cyto - Cytophagaceae, Hyme - Hymenobactereaceae, Coma - Comamonadaceae, Micro - Microbacteriaceae, Meth - Methylobacteriaceae, Rhiz - Rhizobiaceae, Noca - Nocardiaceae, Hyph - Hyphomonadaceae, Mic* - Micrococcaceae, Caul - Caulobacteraceae, Brad - Bradyrhizobiaceae, Osci - Oscillatoriaceae, Xant - Xanthomonadaceae, Chit - Chitinophagaceae, Aura - Aurantimonadaceae.

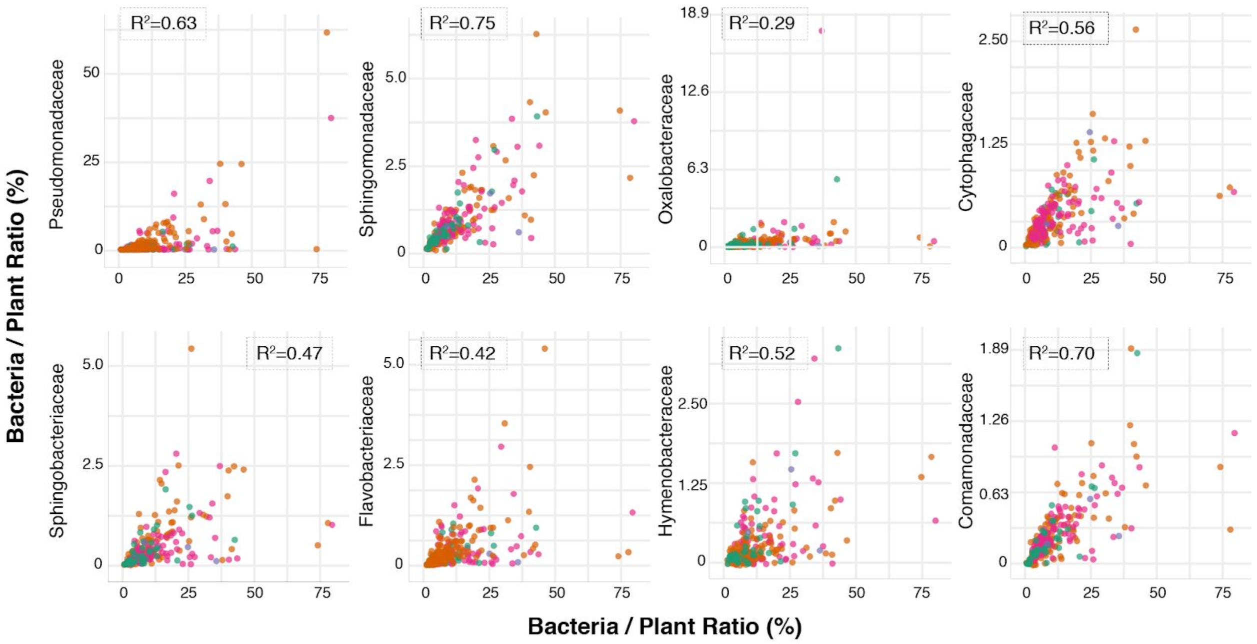


**Supplementary Figure 17: Taxa correlations for bacterial load.**

Scatterplots of individual taxa counts (at family level) and bacterial load, expressed as the fraction of bacteria reads to plant chromosomal reads.

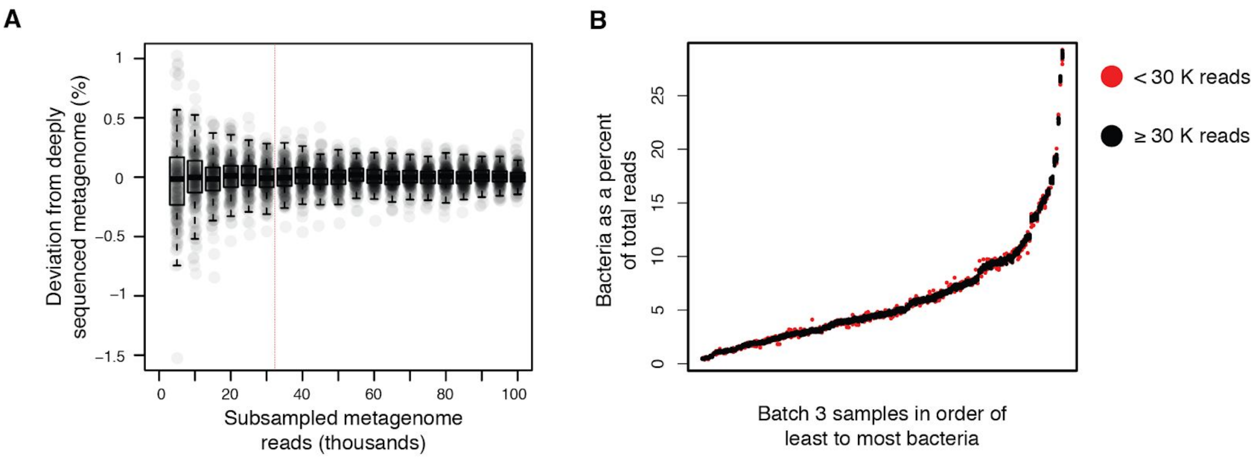


**Supplementary Figure 18. Depth of sequencing to accurately determine microbial load.**

(**A**) In the deep metagenome dataset from Batch 3 plants, bacteria represented between 0.48% and 29% of the total reads in each sample, with a median of 4.6%. To determine how well shallow sequencing could capture these expected percentages of bacteria, each sample in the Batch 3 was rarefied from 5K to 100K reads in increments of 5K. The percent bacteria obtained from each subsample was then subtracted from the percentage of bacteria in the full dataset, and this difference was plotted. With more reads, values tend towards the value obtained by deep sequencing. After 30K reads (to the right of the dotted red vertical line), the total bacterial load estimated for each sample is within 0.5% of the deeply sequenced value. (**B**) Batch 3 samples were ordered from least to most bacteria. The bacterial percentage of total reads as determined by all subsampling efforts was plotted for each sample. The subsamples with fewer than than 30K reads were most responsible for noisy estimates (red points).

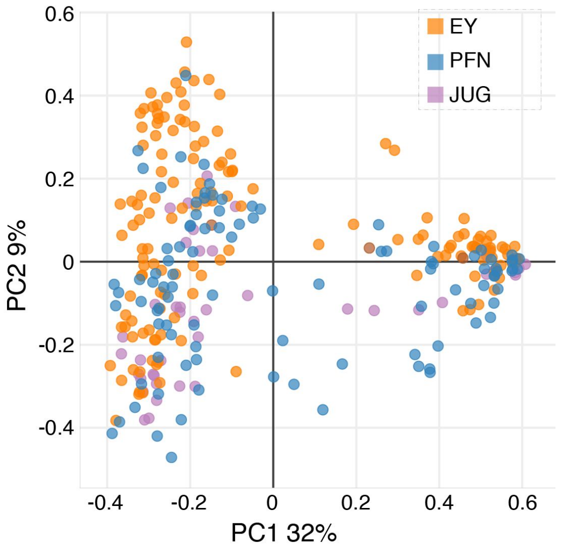


**Supplementary Figure 19: PCA ordination of fungal family abundance.**

Colored by collection sites.
